# The Abl-interactor Abi suppresses the function of the BRAG2 GEF family member Schizo

**DOI:** 10.1101/2020.09.29.317990

**Authors:** Stefanie Lübke, Carina Braukmann, Karl-Heinz Rexer, Lubjinka Cigoja, Susanne F. Önel

## Abstract

Guanine nucleotide exchange factors (GEF) of the BRAG subfamily activate small Arf GTPases, which are pivotal regulators of intracellular membrane traffic and actin dynamics. Here, we demonstrate a novel interaction between the Abl-interactor (Abi) and the BRAG family member Schizo. We mapped the SH3 domain of Abi to interact with the N-terminal region of Schizo. This region is additionally involved in the binding of the cytodomain of the cell adhesion molecule N-cadherin. In *schizo* loss of function mutants, we detected increased amounts of N-cadherin. In contrast, the expression of the GEF (Sec7) and the membrane-binding (pleckstrin homology) domains decreased amounts of N-cadherin, indicating a crucial role of the Sec7-PH module in regulating N-cadherin levels. Unlike other Sec7 GEFs, where the catalytic Sec7 domain is autoinhibited, the Sec7 and PH domain of BRAG2 are constitutively accessible, raising the question how GEF activity is controlled in a spatial and temporal manner. Our genetic analyzes demonstrate that the nature of the Abi Schizo interaction is to antagonize Schizo function and to restore wild-type amounts of N-cadherin.

## Introduction

BRAG proteins are a subgroup of the Arf-guanine nucleotide exchange factor (GEF) family that are mandatory for developmental and physiological processes, e.g. myoblast fusion, neuronal pathfinding and synaptic transmission. However, they also play an important role during disease progression, e.g. cancer metastasis and X-chromosome–linked intellectual disability (D’Souza and Casanova, 2016). The BRAG family is characterized by an N-terminal located calmodulin-binding IQ motif, a catalytic Sec7 domain of ~200 amino acids and a pleckstrin homology (PH) domain that is immediately located downstream of the Sec7 domain. The Sec7 domain stimulates the release of GDP to allow binding of GTP on ADF-ribosylation factor (Arf) family members. These Arf GTPases serve as master regulators of intracellular membrane traffic and actin dynamics. A primary challenge in understanding the activation of small GTPases in development, tissue homeostasis and disease, is to study the molecular mechanism underlying GEF activation.

The human GEFs for Arf family proteins are grouped into six evolutionarily conserved families known as BIG, BRAG/IQSec, Cytohesins, EFA6/PSD, FBX8 and GBFs. Arf GEFs of the Cytohesin and BIG family are regulated by auto-inhibition (DiNitto et al., 2007; Stadler et al., 2011; Richardson et al., 2012). The GEF activity of Cytohesin is suppressed through the Sec7-PH linker and the C-terminal helix/polybasic region that mask the active side in the Sec7 domain (DiNitto et al., 2007). The auto-inhibition of the yeast BIG family member Sec7 depends on an intramolecular interaction that involves the HSD domains (Richardson et al., 2012). Auto-inhibition is released by the binding of membrane-bound Arf-GTP to the PH domain or the HSD1 domain. A positive feedback loop arises through the generation of more Arf-GTP by the Sec7 GEF, which leads to the recruitment of more Arf GEF. In contrast, high resolution crystal structure of the unbound Sec7-PH domain of the Sec7 GEF BRAG2 revealed that the lipid binding side in the PH domain and the Arf-binding site in the Sec7 domain are both constitutively active (Karandur et al., 2017). The finding that the Sec7 and PH domain are constantly accessible for protein-protein and protein-membrane interactions raises the question how the spatial-temporal activation of BRAG2 is achieved.

The *Drosophila* BRAG family member Schizo is required for the guidance of neuronal axons (Önel et al., 2004) and muscle development (Chen et al., 2003, Dottermusch-Heidel et al., 2012). Muscles are multinucleated cells that arise by the fusion of mono-nucleated myoblasts. Despite muscle formation, myoblast fusion is crucial for the maintainance, growth and repair of muscles in mammals and *Drosophila* (Abmayr and Pavlath, 2012; Chaturvedi et al., 2017). However, the precise function of Schizo during myoblast fusion is still unknown.

Rescue experiments with *Drosophila* Arf1, Arf2 and Arf6 that represent the three classes of mammalian Arf GTPases (Donaldson and Jackson, 2011), suggest that Schizo acts through the Arf1-GTPase (Dottermusch-Heidel et al., 2012). In a global yeast two-hybrid screen the cell adhesion molecule N-cadherin was identified as Schizo interaction partner (Dottermusch-Heidel et al., 2012). Genetic interaction studies revealed that the *schizo* myoblast fusion phenotype is suppressed by the loss of N-cadherin. Based on these findings we proposed that the removal of N-cadherin brings the apposing myoblast membranes into close proximity to allow membranes to fuse.

In this study, we have analyzed the function of the N-terminal domain of Schizo. Our data demonstrates that the N-terminal domain is essential for the localization of the Schizo protein to the plasma membrane and the regulation of amounts of N-cadherin. Consistently, the exclusive expression of the Sec7 PH domain of Schizo, which represents an active form of the GEF, reduces amounts of N-cadherin. Finally, we have addressed by which mechanism amounts of N-cadherin are regulated. The finding that Schizo acts through the Arf1-GTPases prompted us to investigate whether CLIC/GEEC endocytosis is involved in the removal of N-cadherin. CLIC/GEEC endocytosis depends on the RhoGAP protein Graf-1. However, neither the deletion of *graf-1* by CRISPR/Cas9 nor the expression of truncated Graf-1 showed increased amounts of N-cadherin as observed in *schizo* mutants. Instead, we found that the Scar/WAVE complex member Abi binds to the N-terminal region of Schizo. Abi, but not Scar/WAVE, antagonizes Schizo function in a dosage-dependent manner. These findings provide a new conceptual framework for the regulation of Schizo activity.

## Results

### The N-terminal region of Schizo is important for protein localization

*schizo* encodes for two Arf GEFs (Schizo P1 and Schizo P2) that only differ in the first 12 amino acids at their N-terminal region and share all the conserved domains (Önel et al., 2004). These Schizo proteins correspond to the Loner isoforms Iso1 and Iso2 that have been described by Chen et al. (2003). In Dottermusch-Heidel et al. (2012) we screened a *Drosophila* yeast two-hybrid cDNA library with the first 753 amino acids of Schizo P2 and identified N-cadherin as interaction partner. To determine whether the N-terminal region of Schizo is important for Schizo P2 localization, we placed the wild-type *schizo* cDNA LP01489 and *schizo* lacking the first 2,3 kb of the *schizo* ORF under the control of UAS activating sequences. The full-length Schizo protein P2 Siz_1-1313_, the Schizo protein Siz_1-753_ from the yeast-two hybrid screen and Siz_753-1313_ lacking the first 2,3 kb of the LP01489 *schizo* cDNA are shown in Fig. 1A. GFP-tagged Siz_1-1313_ and Siz_753-1313_ were expressed in the mesoderm and in somatic muscle cells with the *Mef-*GAL4 driver. In the mesoderm of stage 10 embryos, Siz_1-1313_ and N-cadherin are both present at the plasma membrane (Fig. 1B–B” arrow), but Siz_1-1313_ is also detectable in the cytoplasm (Fig. 1 B–B” asterisks and D–D”). In contrast, Siz_753-1313_ lacking the N-terminal region is only detectable in the cytoplasma and not at the plasma membrane like observed for Siz_1-1313_ (Fig. 1C–C”). In somatic muscles Siz_753-1313_ is expressed in a very specific pattern (Fig. 1E” and E” arrows). This pattern is reminiscent of the Mef2 transcription factor, which is expressed in the nuclei of muscle cells (Bour et al., 1995).

**Figure 1.**
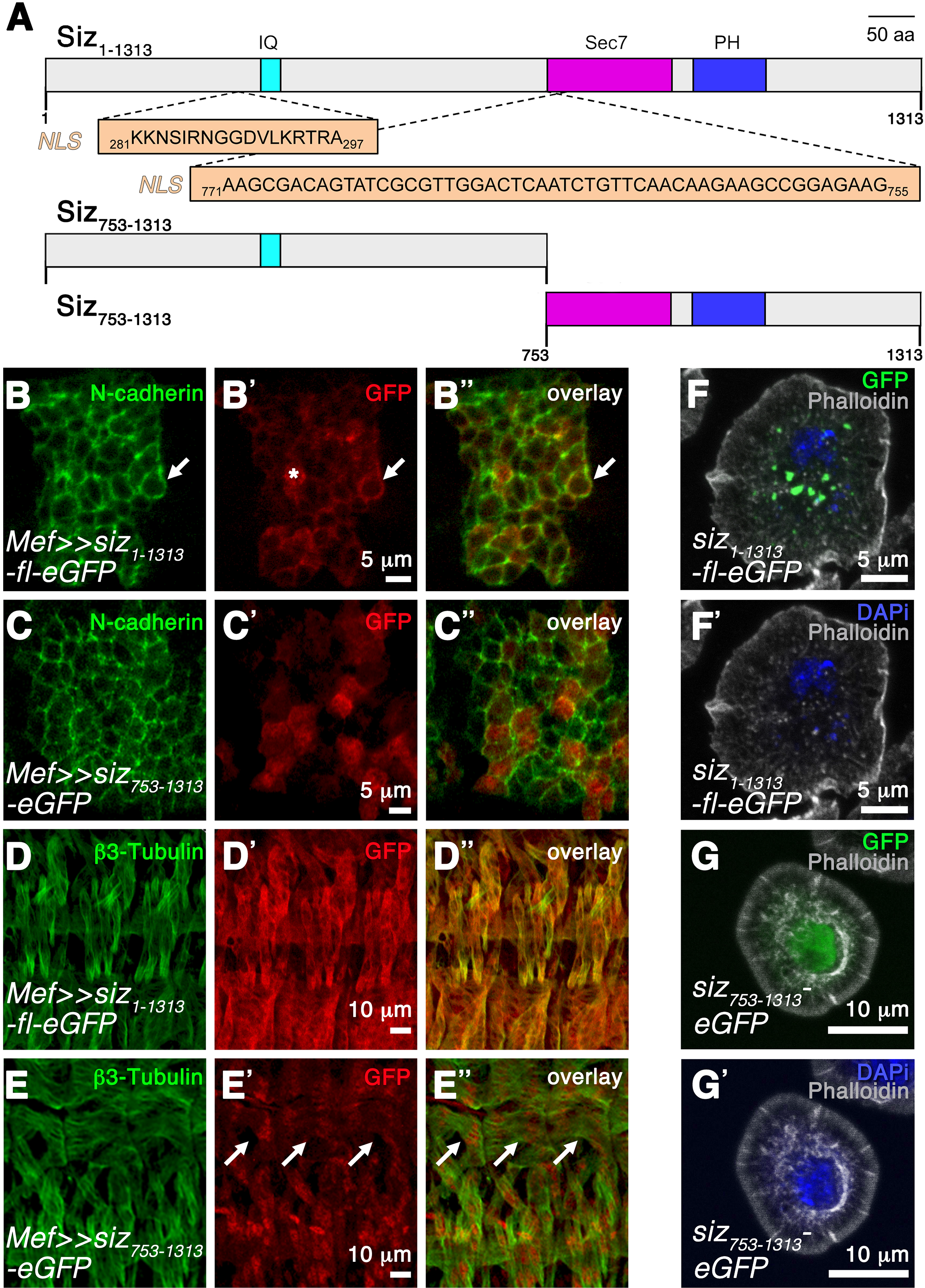
Schizo colocalizes with N-cadherin and undergoes nucleocytoplasmic shuttling in the absence of the N-terminal region. A. Schematic representation of the domain structure of Schizo full-length Siz_1-1313_, Siz_1-753_ and Siz_753-1313_. Caldmodulin domain (IQ), Sec7 domain and pleckstrin homology domain (PH). NLS = nuclear localization signal sequences. B–B”. Higher magnification of a stage 10 embryo expressing Siz_1-1313_-eGFP with Mef-GAL4 in the mesoderm. The embryo was stained with anti-GFP to follow Siz_1-1313_-eGFP expression and with anti-N-cadherin, which is present at the plasma membrane. Siz_1-1313_-eGFP is detectable in the cytoplasm (asterisks) and around the myoblast (arrow). Scale bars 5 μm. C–C”. Higher magnification of a stage 10 embryo expressing Siz_753-1313_-eGFP with Mef-GAL4 in the mesoderm. The embryo was stained with anti-GFP and anti-N-cadherin. In the absence of the N-terminal region, Siz_753-1313_-eGFP is present in the cytoplasma. Scale bars 5 μm. D–D”. Higher magnification of a stage 16 embryo expressing Siz_1-1313_-eGFP with Mef-GAL4 in the mesoderm. The embryo was stained with anti-GFP to follow Siz_1-1313_-eGFP expression and with anti-β3-Tubulin to highlight somatic muscles. Most of the Siz_1-1313_ protein is present in the cytoplasm of muscle cells and at the plasma membrane. Scale bar 10 μm. E–E”. Higher magnification of a stage 16 embryo expressing Siz_753-1313_-eGFP. The embryo was stained with anti-GFP and anti-β3-Tubulin. The cytoplasmic distribution of Siz_753-1313_ in muscle cells seems to be reduced in comparison to Siz_1-1313_. Instead, distinct spots are visible (arrows). F–F’. Transfection of *Drosophila* S2R+ cells with Siz_1-1313_-eGFP. Cells were plated on concanavalin A coated cover slips and stained for DAPi (blue) and Phalloidin (grey). Siz_1-1313_ is distributed in a punctuated manner. Scale bars 5 μm. G–G’. Transfection of *Drosophila* S2R+ cells with Siz_753-1313_-eGFP. Cells were plated on concanavalin A coated cover slips and stained for DAPi (blue) and Phalloidin (grey). In the absence of the N-terminal region Siz_753-1313_ is detectable in the nucleus. Scar bars 10 μm.

To confirm the different localization of Siz_1-1313_ full-length and truncated Siz_753-1313_, both proteins were expressed in *Drosophila* S2R+ Schneider cells (Fig. 1 F, F’, G, G’ and Expanded View Movie EV1). Siz_1-1313_ is distributed in the cytoplasm of S2R+ cells in a punctuated manner (Fig. 1 F and F’). Surprisingly, Siz_753-1313_ is present in the nucleus where it colocalizes with DAPi (Fig. 1G and G’). The import of proteins into the nucleus depends on a short peptide sequences called nuclear localization signal (NLS) sequences. To analyze whether Schizo P2 contains such NLS sequences, we have performed a data bank search as described by Lin and Hu (2013) and identified two predicted NLS sequences in Siz_1-1313_ (Fig. 1A). One of these predicted NLS sequences is located in the Sec7 domain of Schizo P2 and is still present in Siz_753-1313_. However, the expression of Siz_753-1313_ in the mesoderm of wild-type embryos and its nucleocytoplasmic shuttling does not disturb myoblast fusion. Taken together, these results show that the N-terminal region of Schizo P2 is essential for the localization of the Schizo protein to the plasma membrane. In the absence of this region Schizo translocate to the nucleus although the Sec7 and PH domain that mediate lipid binding are still present.

### N-cadherin amounts are increased in homozygous *schizo* mutant embryos

Schizo function is required in the two types of *Drosophila* myoblasts: founder cells (FCs) and fusion-competent myoblasts (FCMs) (Dottermusch-Heidel et al., 2012), where it interacts with N-cadherin (Fig. 2A). The loss of Schizo function leads to severe defects in myoblast fusion (Fig. 2A’ and A”). In the CNS, *schizo* mutants lack commissural axons (Fig. 2 B’ and B”, arrows) due to increased levels of the axon guidance molecule Slit (Önel et al., 2004). To assess the importance of the Sec7 and PH domain for Schizo P2 function, we generated Siz-ΔSec7 and Siz-ΔPH deletion mutants. Expression of Siz_1-1313_ in the mesoderm with *twist*-GAL4 rescues the myoblast fusion phenotype (Fig. 2E), whereas homozygous mutants expressing Siz-ΔSec7 or Siz-ΔPH in their mesoderm still showed a myoblast fusion phenotype (Fig. 2F and G). These data confirm that the Sec7 and PH domains are both pivotal for Schizo function.

**Figure 2.**
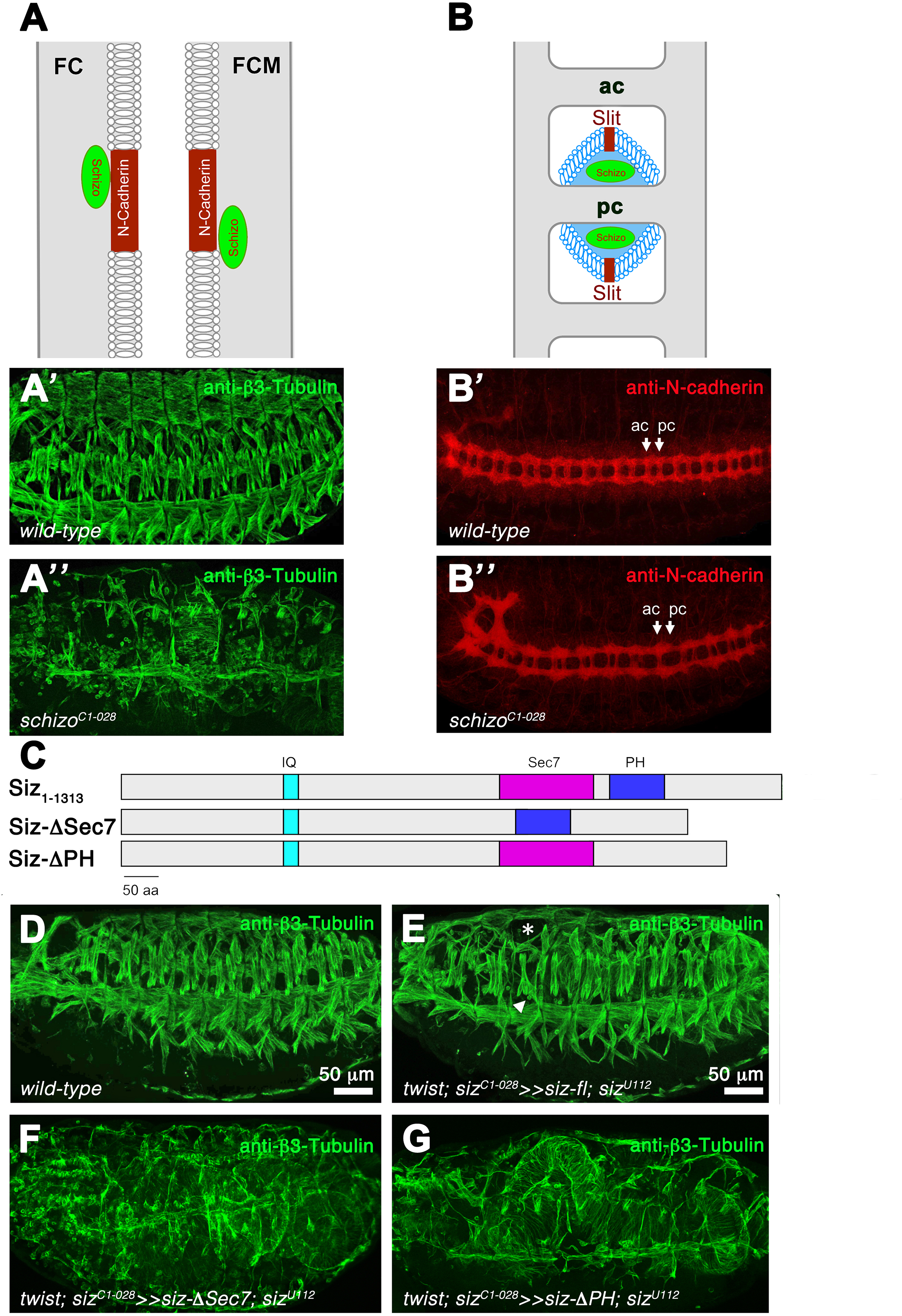
The Sec7 and PH domains are important for Schizo function. A. Schematic representation of N-cadherin and Schizo expression in myoblasts. In *Drosophila*, myoblasts are subdivided into founder cells (FCs) and fusion-competent myoblasts (FCMs). N-cadherin and Schizo are expressed in both myoblast-types, and the N-terminal region of Schizo binds to the intracellular domain of N-cadherin. A’. Lateral view of a stage 16 wild-type embryo stained with anti-β3-Tubulin to visualize somatic muscles. At the end of embryogenesis, a repeated pattern of multinucleated muscles per hemisegment is visible. A”. Lateral view of a stage 16 embryo homozygous for the *schizo^C1-028^* allele. The embryo was stained with anti-β3-Tubulin. The multinucleated muscle pattern is disturbed in *schizo* mutants, since myoblasts fail to fuse. B. Schematic representation of the *Drosophila* ventral nerve cord. Neurons send out their axons towards the midline to form the anterior (ac) and posterior (pc) commissures. To ensure that commissural axons only cross the midline once, the midline glia cells (blue) secrete the repellent Slit. Schizo is expressed in midline glia cells and antagonizes Slit signalling. B’. Ventral view of a stage 16 wild-type embryo stained with anti-N-cadherin showing the ladder-like central nervous system. ac = anterior commissure, pc = posterior commissure. B”. Ventral view of a stage 16 homozygous *schizo^C1-028^* mutant embryo stained with anti-N-cadherin. The posterior commissures (pc) are predominantly missing in the hypomorphic *schizo* allele *schizo^C1-028^*. C. Schematic diagrams of the domain organization in Siz_1-1313_, SizΔSec7 and SizΔPH. D. Lateral view of stage 16 wild-type control embryo stained with anti-β3-Tubulin. E. Transheterozygous *schizo^C1-028^/schizo^U112^* mutant embryo expressing Siz_1-1313_ in the mesoderm with *twist*-GAL4. The expression of Siz_1-1313_ rescues the *schizo* myoblast fusion phenotype. F. Lateral view of stage 16 embryo stained with anti-β3-Tubulin. Transheterozygous *schizo^C1-028^/schizo^U112^* mutant embryo expressing SizΔSec7. In the absence of the Sec7 domain, Siz fails to rescue the *schizo* mutant phenotype. G. Lateral view of stage 16 embryo stained with anti-β3-Tubulin. Transheterozygous *schizo^C1-028^/schizo^U112^* mutant embryo expressing SizΔPH. In the absence of the PH domain, Siz fails to rescue the *schizo* mutant phenotype.

However, the question remained whether the Sec7-PH module is crucial for regulating amounts of N-cadherin. To address this question, we first elevated the amount of N-cadherin in homozygous *schizo* mutants and in embryos expressing dominant-negative Arf1 (ArfT31N) by using two different approaches. Consistent with our previous hypothesis that was based on genetic interaction studies, we observed increased amounts of N-cadherin in whole-mount embryos (Fig. 3A’, A”, B’, B”, C’ and C”). To analyze the amount of N-cadherin, we used cytofluorograms of single embryos (Fig. 3A”, B” and C”) and measured the fluorescence intensity from 6 to 15 embryos (Fig. 3F). The activity of the Schizo homologue BRAG2 that only possesses the Sec7-PH module has been shown to possess a 10-fold higher activity towards Arf1 and a 15–20-fold higher activity in the presence of PIP_2_ (Jian et al., 2012; Aizel et al., 2013). Conversely, to investigate whether an increased activity of Schizo is able to decrease amounts of N-cadherin in whole mount-embryos, we generated Siz-Sec7-PH_753-1081_ transgenic flies (Fig. 3D). The expression of Siz-Sec7-PH_753-1081_ in muscles with *Mef*-GAL4 leads to severe defects in myoblast fusion (Fig. 3E, E’). Furthermore, we observed a punctated distribution of Siz-Sec7-PH_753-1081_ in myoblasts, which was not observed with Siz_1-1313_ (Fig. 3E”; Fig. 1B–B” and D–D”). Additionally, we examined the N-cadherin fluorescence intensity of embryos expressing Sec7-PH_753-1081_ in the mesoderm with *twist*-GAL4 or with *Mef*-GAL4 in muscle cells. With both GAL4 drivers, we observed decreased amount of N-cadherin (Fig. 3G). These data show that the Schizo full-length protein is required for the removal of N-cadherin and that Schizo-Sec7-PH_753-1081_ represents an activated form of Schizo.

**Figure 3.**
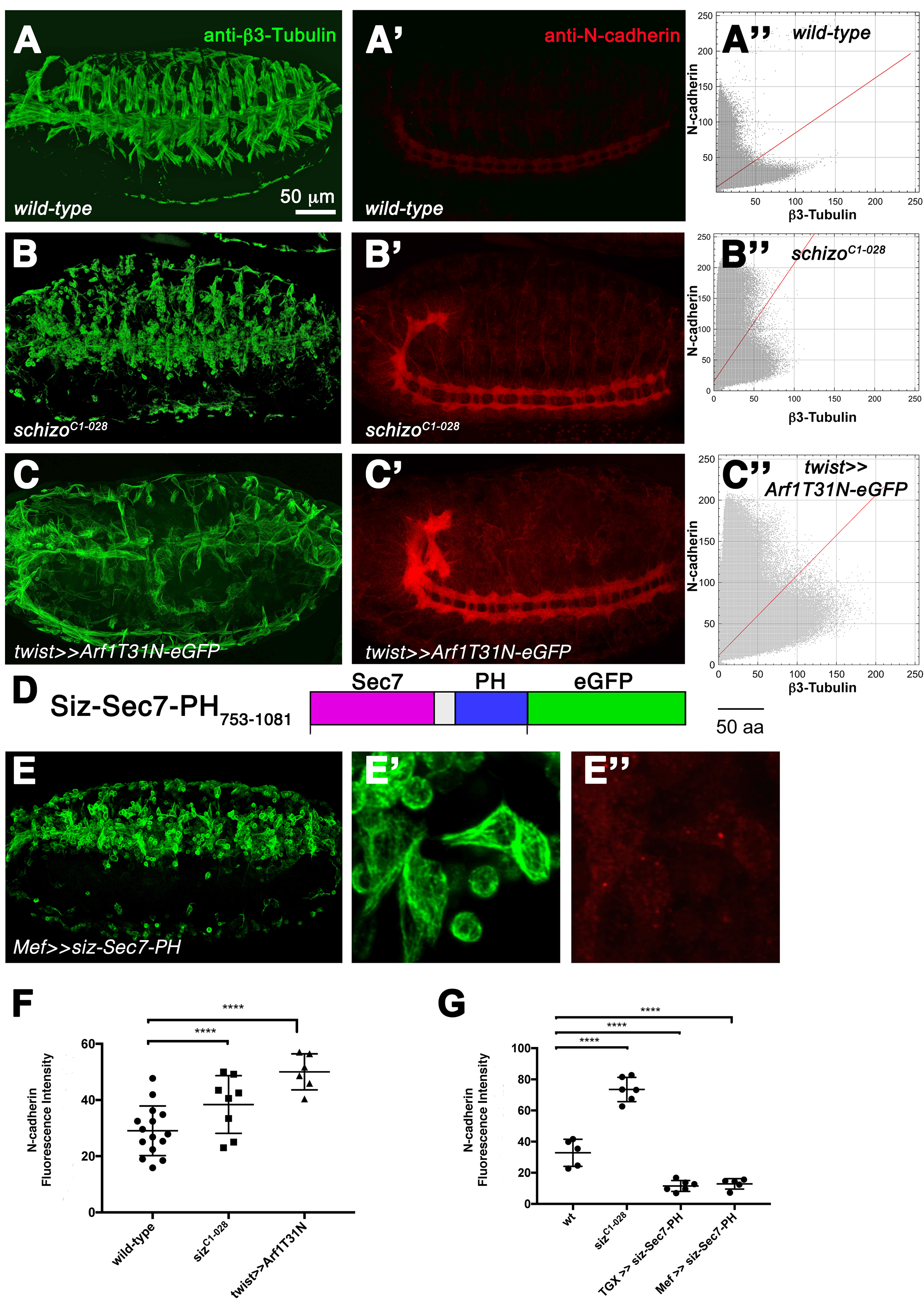
Amounts of N-cadherin are decreased in *schizo* mutants. A,A’. Wild-type control. Lateral view of a stage 16 embryo stained with anti-β3-Tubulin (A) and anti-N-cadherin (A’). A”. Cytofluorogram of the embryo shown in A and A” comparing the fluorescence concentration of β3-Tubulin and N-cadherin in the wild-type. B,B’. Lateral view of a stage 16 homozygous *schizo^C1-028^* mutant embryo stained with anti-β3-Tubulin (B) and anti-N-cadherin (B’). B”. Cytofluorogram of the embryo shown in B and B” comparing the fluorescence concentration of β3-Tubulin and N-cadherin in a homozygous *schizo^C1-028^* mutant embryo. Amounts of N-cadherin are increased in homozygous *schizo^C1-028^* mutants. C,C’. Expression of UAS-*Arf1T31N-eGFP* with *twist*-GAL4 in the background of a wild-type embryo. Lateral view of a stage 16 embryo stained with anti-β3-Tubulin (C) and anti-N-cadherin (C’). The muscle pattern is disturbed in *twist*-GAL4≫UAS-*Arf1T31N-eGFP* expressing embryos. C”. Cytofluorogram of the embryo shown in C and C” comparing the fluorescence concentration of β3-Tubulin and N-cadherin. Amounts of N-cadherin are increased in the embryo expressing *twist*-GAL4≫UAS-*Arf1T31N-eGFP*. D. Schematic representation of Schizo containing only the Sec7-PH domain, which was used for overexpression studies in E–E” and G. E–E”. The expression of UAS-*siz-Sec7-PH*-eGFP in the mesoderm of wild-type embryos with *Mef*-GAL4 induces severe defects in myoblast fusion (E and E’). The muscles and unfused myoblasts of a stage 16 embryo are visualized by anti-β3-Tubulin. An anti-GFP staining revealed that Siz-Sec7-PH is distributed in a punctuated manner in myoblasts (E”). F. Quantification of the amounts of N-cadherin in wild-type, homozygous *schizo^C1-028^* mutant embryos and embryos expressing UAS-*Arf1T31N-eGFP* with *twist*-GAL4. The total fluorescence intensity of 6 to 15 embryos was measured for each experiment. Bars represent mean ± s.d. P-values were calculated using the Dunnett’s multiple comparison test. p^***^< 0,0001 compared to wild-type. In *Mef*-GAL4≫UAS-*Arf1T31N-eGFP* expressing embryos increased amounts of N-cadherin were not observed. G. Quantification of the amounts of N-cadherin in wild-type, homozygous *schizo^C1-028^* mutant embryos, embryos expressing UAS-*siz-Sec7-PH-eGFP* with *Mef-*GAL4 and *twist*-GAL4. The total fluorescence intensity of 5 to 6 embryos was measured for each experiment. In *Mef*-GAL4≫UAS-*siz-Sec7-PH-eGFP* and *twist*-GAL4≫UAS-*siz-Sec7-PH-eGFP* embryos amounts of N-cadherin are decreased. Bars represent mean ± s.d. P-values were calculated using the Dunnett’s multiple comparison test. p^***^< 0,0001 compared to wild-type.

### The Scar/WAVE complex member Abi interacts physically with the RhoGAP protein Graf-1 and Schizo

The finding that amounts of N-cadherin are increased in homozygous *schizo* mutants and the punctuated distribution of Schizo-Sec7-PH_753-1081_ in muscles, prompted us to investigate whether Schizo controls N-cadherin amounts by endocytosis. The observation that the *schizo* mutant phenotype can be rescued by GTP-bound Arf1 suggests that Schizo acts via Arf1 (Dottermusch-Heidel et al., 2012). Consistently, we detect a transient colocalization between Siz_1-1313_ and Arf1 in *Drosophila* S2R+ cells (Appendix Fig. S1). The small Arf1-GTPase is involved in the endocytosis of lipid-anchored proteins such as GPI-APs and does not involve Dynamin. The endocytotic structures are termed GEECs (GPI-AP enriched early endosomal compartments). The molecular mechanism of this pathway is initiated by the recruitment of the Sec7 GEF GBF1, which activates Arf1 (Gupta et al., 2009). Arf1 recruits the RhoGAP protein Graf-2 (alias ARHGAP10, alias ARHGAP21) to the cell surface (Kumari and Mayor, 2008). The activity of this protein complex promotes GTP-hydrolysis on Cdc42, which is necessary for the endocytosis process. Additionally, Graf-1 (alias ARHGAP26) has been described to colocalize with activated Cdc42 and controls like Graf-2 Cdc42 activity (Lundmark et al., 2008). During muscle development, the down-regulation of Graf-1 in murine C2C12 cells or the loss of Graf-1 or Graf-2 in primary myoblasts significantly reduces the capability of myoblasts to fuse (Doherty et al., 2011; Lenhart et al., 2014). Due to the role of Graf in GEEC endocytosis and its role in mammalian myoblast fusion, we examined Graf function in *Drosophila*.

*Drosophila* Graf-1 contains like the mammalian Graf proteins a BAR, PH, RhoGAP and SH3 domain (Kim et al., 2017) (Fig. EV1A). To determine whether *Drosophila* Graf-1 is involved in the removal of N-cadherin, we first performed protein interaction studies between Graf-1 and Schizo and the intracellular domain of N-cadherin. Although we did not detect any interaction between Graf-1 full-length and N-cadherin, we found that Graf-1 lacking the BAR domain interacted with the intracellular domain of N-cadherin (Fig. EV1A, Table 1, Appendix Fig. S2). The BAR domain of mammalian Graf-1 has been shown to directly interact with the GAP domain to inhibit its activity (Eberth et al., 2009). We assume that the failed interaction between Graf-1 full-length and the intracellular domain of N-cadherin is caused by the autoinhibited conformation of Graf-1. Domain mapping experiments revealed that the SH3 domain of Graf-1 is responsible for the interaction of GrafΔBAR with the intracellular domain of N-cadherin (Fig. EV1A, Appendix Fig. S3). However, we found no protein interactions between Graf-1 and Schizo in the yeast-two hybrid assay. Graf-1-eGFP is distributed in a punctuated manner in *Drosophila* S2R+ cells (Fig. EV1B and C) and partially colocalizes with Schizo and GTP-bound Arf1 (Fig. EV1D and E). To determine whether Graf is involved in the removal of N-cadherin, we utilized the CRISPR/Cas9 strategy to generate *graf* deletion mutants (Fig. EV1F). Although, we identified small deletions in the BAR domain of *graf*, homozygous *graf* mutants were viable and showed no defects in myoblast fusion (Fig. EV1G and H). In addition, we quantified amounts of N-cadherin in homozygous *graf* mutants and overexpressed UAS-*grafΔBAR* and UAS-*grafDBARΔSH3* with *Mef*-GAL4 in wildtype embryos (Fig. EVA1 I). However, we failed to observe increased amounts of N-cadherin. Based on these findings, we propose that Graf-1-dependent endocytosis is not essential for N-cadherin removal.

**Table 1.**
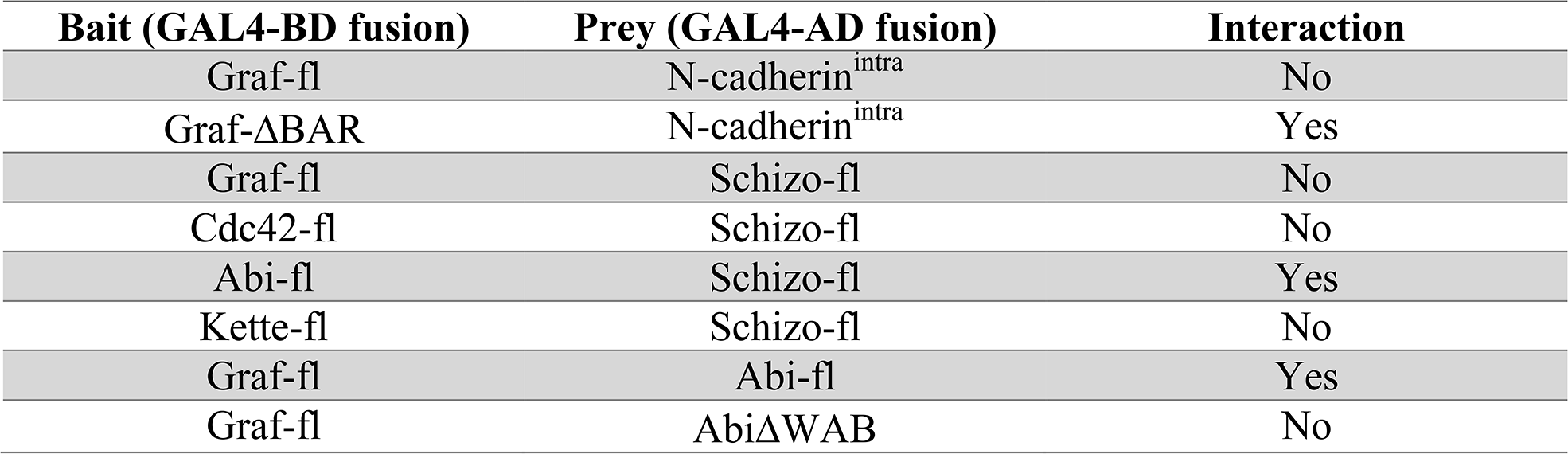
Yeast-two hybrid analysis of Graf-1, Cdc42, Abi and Kette interactions as indicated by growth of media (SD/-Leu/-Trp/-His/-Ade/X-α-Gal).

The actin polymerization machinery has been implicated to aid in clathrin-independent endocytosis (Chadda et al., 2007; Römer et al., 2010; Sathe et al., 2018). Indeed, actin polymerization seems to be a key step to power local membrane deformation and carrier budding in clathrin-independent endocytosis (Hinze and Boucrot, 2018). Since Arp2/3- and Formin-dependent F-actin polymerization is essential for the fusion of myoblasts (Kim et al., 2007; Massarwa et al., 2007; Richardson et al., 2007; Schäfer et al., 2007; Berger et al., 2008; Deng et al., 2015), we determined whether members of the Arp2/3 activation machinery or the formin Diaphanous interact with Schizo (Table 1). Surprisingly, we identified Abi, a component of the Scar/WAVE complex, to interact with Schizo and Graf-1 (Table 1, Fig. 4 and Appendix Fig. 4).

**Figure 4.**
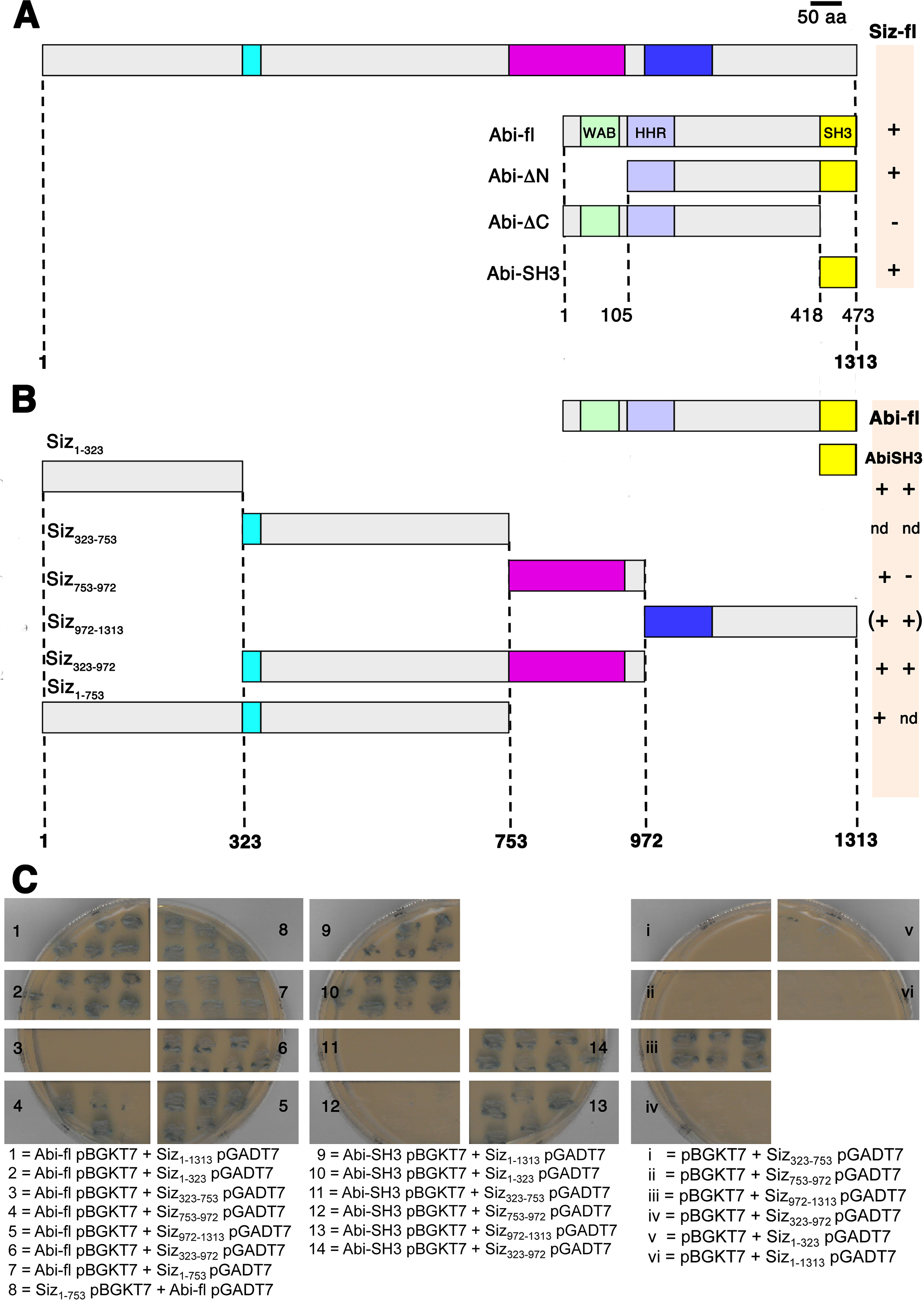
Yeast two-hybrid tests of the binding of Abi and the Abi SH3 domain to the N-terminal domain of Schizo. A. Schematic representation of Schizo and Abi deletions employed to study protein interactions. B. Schematic representation of Abi and Schizo deletions employed to study protein interactions. C. Yeast two hybrid tests. As indicators for interaction the growth on SD-Ade/-Leu/-His/-Trp plates and the implementation of α-Galaktosidase by the Galactosidase MEL1 were used. Plates were incubated for 48 h. − no growth and blue coluor in 48 h; + growth and blue coluor in 48 h. In 1 to 7 Abi full-length cloned into the bait vector pBKGT7 from Clontech was tested for protein interaction with Schizo deletions cloned into the prey vector pGADT7. Schizo full-length Siz_1-1313_, Siz_1-323_, Siz_753-972_, Siz_972-1313_ and Siz_323-972_ showed an interaction with Abi. Control experiments (i–vi) revealed an interaction of Siz_972-1313_ pGADT7 with the empty pBGKT7. In 8 the N-terminal region of Siz_1-753_ cloned into the pBGKT7 interacted with Abi full-length cloned into the pGADT7. In 9 to 14 only the SH3 domain of Abi into the pBGKT7 was used for interaction studies with Schizo deletions. An interaction with Siz_1-1313_, Siz_1-323_, Siz_972-1313_ and Siz_323-972_ was observed.

### The SH3 domain of Abi interacts with the N-terminal region of Schizo

We next mapped the interaction domain on Abi by testing deletion mutants of Abi for binding to full-length Siz_1-1313_. Full-length Abi and the deletion mutant proteins examined are depicted schematically in Figure 4A. The deletion of the carboxy-terminal SH3 domain eliminated the interaction with Schizo Siz_1-1313_ (Fig. 4A). Subsequently, we performed the converse experiment and only used the SH3 domain of Abi for determining the interaction with Schizo (Fig. 4A and C). These data indicate that the SH3 domain is mandatory for the observed interaction. To identify the interaction domain on Schizo, we utilized the deletion mutant proteins illustrated in Figure 4B. Abi full-length interacts with Siz_1-323_, Siz_753-972_, Siz_972-1313_ and Siz_1-753_. As demonstrated in the controls, Siz_972-1313_ shows a false positive interaction with the empty pBGKT7 vector. By using the SH3 domain of Abi as bait protein, we could confirm the interaction of Siz_1-323_, Siz_972-1313_ and Siz_1-753_, but not with Siz_753-972_. From these data, we concluded that the N-terminal region of Schizo from 1–753 amino acid is responsible for interacting with Abi. Because of the observed protein interaction, we next transfected *Drosophila* S2R+ cells with UAS-*siz-mcherry* and UAS-*abi-eGFP* to determine the distribution of both proteins in Schneider cells. We found that UAS-*abi-eGFP* colocalizes with full-length UAS-*schizo-mcherry* and that both proteins are distributed in a punctuated manner (Appendix Fig. S5 and Expanded View Movie EV2).

### Abi and Schizo serve antagonistic functions

The observation that Abi and Schizo colocalizes in a punctuated manner raises the question whether Abi and Schizo act in concert to regulate amounts of N-cadherin by endocytosis. To determine whether both proteins contribute to the same process, we performed epistasis experiments and used meitotic recombination to generate *schizo abi* double mutants. In a first attempt we analyzed the muscle phenotype of the hypomorphic *schizo* allele *siz^C1-028^*, the *abi* null allele *abi^Δ20^* and *siz^C1-028^ abi^Δ20^* double mutants (Fig. 5Aa–e). We observed that homozygous *siz^C1-028^ abi^Δ20^* double mutants show a strong myoblast fusion phenotype like *siz^C1-028^* (Fig. 5Ad and Ab). The muscle phenotype of *abi^Δ20^* null mutants is comparable to the wild-type muscle pattern (Fig. 5Ac and Aa), but some muscles are missing. If *schizo* and *abi* both contribute in regulating amounts of N-cadherin, we expected to enhance the *abi* muscle phenotype by reducing the *schizo* gene dose. However, the musculature of transheterozygous *abi^Δ20^/Df(abi)* mutants lacking one copy of *schizo* look like the musculature of homozygous *abi^Δ20^* mutant embryos (Fig. Ae and Ac). From the myoblast fusion phenotype of the epistasis experiments we cannot conclude whether *schizo* and *abi* contribute together in regulating amounts of N-cadherin. To verifty, we determined amounts of N-cadherin in homozygous *siz^C1-028^*, *abi^Δ20^/Df(abi)* single and *siz^C1-028^ abi^Δ20^* double mutants. Figure 5Af shows the measurement of the N-cadherin fluorescence intensity from 5 to 12 embryos. Interestingly, we found that amounts of N-cadherin are reduced to wild-type levels in homozygous *siz^C1-028^ abi^Δ20^* double mutant embryos. The reduced amounts of N-cadherin in the double mutants indicate that *schizo* and *abi* might act antagonistically. Further support for this notion comes from the analyses of the central nervous phenotype with anti-N-cadherin (Fig. 5Ba–f). N-cadherin is not only expressed in the mesoderm and during myoblast fusion, but also shows a strong expression in the central nervous cord (Iwai et al., 1997). When we imaged *siz^C1-028^ abi^Δ20^* double mutants to determine the fluorescence intensity of N-cadherin, we noticed that the commissural phenotype of homozygous *schizo* mutants (Fig. 5Bb arrowheads) is suppressed in some hemisegments of *siz^C1-028^ abi^Δ20^* double mutants (Fig. 5Bd arrowheads).

**Figure 5.**
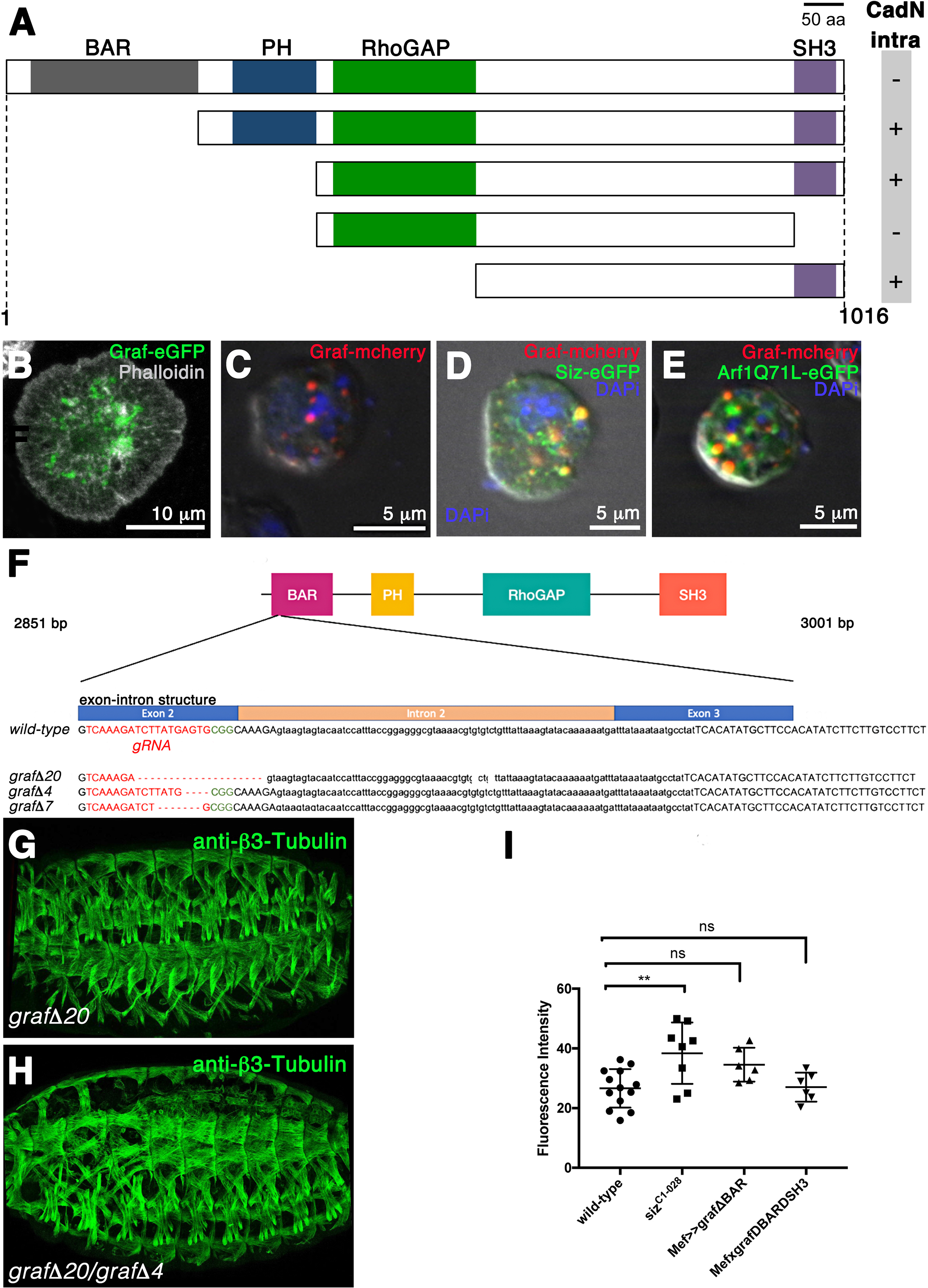
Abi and Schizo serve antagonistic functions. A. *Drosophila* stage 16 embryos stained with anti-β3-Tubulin. Anterior is left, lateral view. The muscle patterns of (a) a wild-type control, (b) a homozygous *siz^C1-028^*, (c) a transheterozygous *abi^Δ20^*/Df(*abi*), (d) a homozygous *abi^Δ20^ siz^C1-028^* and (e) a transheterozygous *abi^Δ20^*/Df(*abi*) mutant lacking on copy of *siz^C1-028^* is shown. The muscle pattern of the homozygous *abi^Δ20^ siz^C1-028^* mutant embryo looks like the muscle pattern of homozygous *siz^C1-028^* mutants. The reduction of *schizo* in *abi^Δ20^*/Df(*abi*) mutants did not enhance the muscle phenotype of *abi^Δ20^*/Df(*abi*) mutants. (f) Quantification of amounts of N-cadherin in wild-type, homozygous *siz^C1-028^* and *abi^Δ20^* mutants and homozygous *siz^C1-028^ abi^Δ20^* double. The total fluorescence intensity of 5 to 12 embryos was measured for each experiment. Bars represent mean ± s.d. P-values were calculated using the Dunnett’s multiple comparison test. P^***^< 0,0006 compared to wild-type; n.s. = not significant. B. Ventral view of stage 16 embryos. The commissural phenotype of wild-type, homozygous *siz^C1-028^*, transheterozygous *abi^Δ20^*/Df(*abi*) and homozygous *abi^Δ20^ siz^C1-028^* double mutant embryos was examined with anti-N-cadherin (red). (a) Wild-type embryo. (b) The anterior commissure is missing in a homozygous siz^C1-028^ mutant embryo (arrow). (c) Transheterozygous *abi^Δ20^*/Df(*abi*) mutant embryos display A wild-type CNS. (d) In homozygous *abi^Δ20^ siz^C1-028^* double mutants the anterior and posterior commissures are axons abnormally cross the midline (arrows). Abi is a member of the Scar/WAVE complex. (e) The abnormal crossing of axons can be also observed in homozygous *scar^k13811^* mutants (arrow). (f) Homozygous *scar^k13811^ siz^C1-028^* double mutant embryos show an enhancement of the *schizo* mutant phenotype. Anterior and posterior commissural axons fail to cross the midline. C. Modification of the Schizo overexpression phenotype. Scanning electron micrographs of adult eyes. (a) The overexpression of UAS-*schizo* in photoreceptor cells with *GMR*-GAL4 induces a rough eye phenotype (*GMR*-GAL4/+; UAS-*siz*/+). (b) This phenotype can be suppressed by taking out one copy of *abi^Δ20^* (*GMR*-GAL4/+; UAS-*siz*/ *abi^Δ20^*). (c) Taking out one copy of *scar^k13811^* does not suppress the *GMR*-GAL4/+≫UAS*-siz/+*-induced phenotype.

Abi is a member of the Scar/WAVE complex, which is required for the activation of the Arp2/3 complex that nucleates branched F-actin filaments (Rotty et al., 2014). The nucleation ability of the Arp2/3 complex depends on nucleation promoting factors of the WASp family to which Scar/WAVE belongs to (Tyler et al., 2016). Besides its function as member of the Scar/WAVE complex, Abi is also known to interact physically with WASp (Bogdan and Klämbt, 2003). The finding that *abi* antagonizes *schizo* during myoblast fusion and commissural formation suggests that also other members of the Scar/WAVE complex, *scar/wave* counteract *schizo*. To test this assumption, we generated *scar siz* double mutants by using the hypomorphic *scar* allele *scar^k13811^*. Commissures are reduced in the *siz* alleles *siz^C1-028^* (Fig. 2A, Fig. 5Bb arrowheads). In homozygous *scar^k13811^* mutants, commissures are not reduced, but both commissures are sometimes observed in close proximity (Fig. 5Be arrowheads). In contrast, we found that the commissural phenotype of homozygous *siz^C1-028^* mutant embryos is clearly enhanced in homozygous *scar^k13811^ siz^C1-028^* double mutant embryos (Fig. 5Bf, arrowheads). Taken together, these data suggest that only *abi* counteracts *schizo*, but not *scar/wave*.

In *Drosophila* about 60% of the genome is maternally contributed as mRNA to ensure embryonic development (De Renzis et al., 2007; Lecuyer et al., 2007). The mRNA of *abi* and *scar* are maternally transcribed (Zallen et al., 2002; Lin et al., 2009). We found that the *schizo* mRNA is still detectable in the *schizo* deficiency Df(3L)*ME178* at embryonic stage 11 when myoblasts start to fuse and at stage 14 (Appendix Fig. 6). This suggests that during embryogenesis reduced protein levels of Abi are required to antagonize Schizo function, but not Scar/WAVE. To support this notion, we overexpressed Schizo in photoreceptor cells in adult flies that have no maternal mRNA using the eye-specific *GMR*-GAL4 driver line. Expression of UAS-*siz* leads to a rough-eye phenotype (Fig. 5Cb). This rough-eye phenotype is suppressed in flies heterozygous for the *abi^Δ20^* mutation (Fig. 5Cc). Consistently with our previous results, the rough-eye phenotype is not suppressed in flies heterozygous for the hypomorphic *scar* allele *scar^k13811^* (Fig. 5Cd). In addition, we observed no suppression of the rough-eye phenotype in flies heterozygous for the *rac* null allele *rac1^J11^*, which is another member of the Scar/WAVE complex (Appendix Fig. 7B). Moreover, we examined whether the Abi interacting partner WASp is able to antagonize Schizo function. However, we found that the reduction of the *wasp* dosage using the dominant-negative EMS allele *wasp^3D3-035^* enhances the *GMR-*GAL4≫UAS-*siz* induced rough-eye phenotype (Appendix Fig. 7C). In summary, these experiments confirm our hypothesis that only reduced levels of Abi antagonize Schizo function.

## Discussion

Understanding the multiple regulatory layers of GEF activation that allows coordinating the GDP/GTP exchange on small GTPases is an important issue. To date, the multitude of molecular interactions leading to GEF activation are most advanced for the Ras activator Son of Sevenless (Bandaru et al., 2019). Studies on the subfamily of Arf GEFs that carry a PH domain associated with a catalytic Sec7 domain have concentrated on the interaction of the GEF with Arf GTPases and phospholipids. Membrane recruitment of Arf GTPases is mediated by a myristoylated N-terminal amphipathic helix (Franco et al., 1995, Goldberg, 1998; Liu et al., 2009) and is crucial for the activation by the GEF (Pasqualato et al., 2001; Randazzo et al., 1995). The PH domain binds phophatidyl inositol 3,4,5-triphosphate (PIP_3_) and phosphatidyl insositol 4,5-bisphosphate (PIP_2_) (Chardin et al., 1996; Kavran et al., 1998; Karlund et al., 2000). In structural studies with ^myr^Arf/BRAG2 bound to a PIP_2_-containing bilayer, the myristoylated N-terminal helix of Arf is close to the Sec7 domain and it has been proposed that the Sec7 domain might recognize conformational information from the amphipathic helix (Karandur et al., 2017). However, so far the influence of receptor binding for Arf GEF activation has not been taken into account.

In this study, we found that N-cadherin levels were elevated in mutants of the Arf1 GEF *schizo*. These findings are in line with studies on mammalian GEP_100_/BRAG2. The siRNA-mediated depletion of GEP_100_/BRAG2 in HepG2 cells resulted in increased E-cadherin content (Hiroi et al., 2006). Furthermore, increased amounts of β1-integrin were observed in the “knock-down” of BRAG2 in HeLa cells and it was proposed that BRAG2 served specifically for β1-integrin internalization (Dunphy et al., 2006).

The finding that the member of the BRAG2 subfamily Schizo interacts with N-cadherin via its N-terminal region let us investigate the importance of this region for Schizo function. Our data imply that the N-terminal region provides additional layers of GEF regulation. First, we identified that Schizo undergoes nucleocytoplasmic shuttling in the absence of the N-terminal region. Mammalian BRAG2a and BRAG2b were detected in the nuclei in HeLa and MDCK cells when overexpressed. Moreover, after treatment with leptomycin B, which inhibits the Crm1/exportin1 nuclear-export machinery, both proteins were exclusively found in the nucleus (Dunphy et al., 2006). BRAG2 possesses like Schizo a nuclear localization sequence in the Sec7 domain. However, whether the nucleocytoplasmic shuttling of Schizo is part of a regulatory pathway needs to be further investigated. Second, we found that Abi binds to the N-terminal region of Schizo. Abi has been reported to regulate actin polymerization by formation of complexes with Scar/WAVE and WASp (Innocenti et al, 2005). Furthermore, it has been described to modulate EGFR endocytosis (Tanos and Pendergast, 2007). Besides, Abi-1 has been identified to interact with the Ras activator Son of Sevenless (Scita et al., 1999; Fan and Goff, 2000).

Vesicle scission in Clathrin-independent endocytosis depends on specialized actin-based platforms (Doherty and McMahon, 2009; Mayor et al., 2014). However, there is no unifying theme and multiple mechanisms may co-exist. The molecular machinery of CLIC/GEEC-dependent endocytosis involves the activation of Arf1 by the Arf GEF GBF1 and the RhoGAP protein Graf that removes Cdc42 from the plasma membrane (Kumari and Mayor, 2008; Gupta et al., 2009). A recent study has addressed the spatio-temporal localization of known molecules affecting CLIC/GEEC endocytosis by using by using real-time TIRF microscopy (Sathe et al., 2018). In this study it was reported that Arp3 recruitment occurred earlier to endocytic vesicles than Cdc42. Furthermore, N-WASp failed to recruit to form CLIC/GEEC endocytotic sites. These data imply alternative pathways for Arp2/3 activation in CLIC/GEEC endocytosis. In *Drosophila*, WASp lacking the Cdc42-binding domain (WaspΔCRIB) is still able to rescue the adult phenotype of *wasp* mutants and it was proposed that other elements than Cdc42 contribute to *Drosophila* WASp activation (Tal et al., 2002). Such an alternative element might be Abi. However, our genetic interaction studies suggested that Abi counteracts Schizo function.

Dissecting the function of Arp2/3-dependent endocytosis during myoblast fusion is challenging since the fusion of myoblasts depends on Scar/WAVE- and WASp-dependent Arp2/3 activation (Kim et al., 2007; Massarwa et al., 2007; Richardson et al., 2007; Schäfer et al., 2007; Berger et al., 2008). This might explain why the myoblast fusion phenotype in *abi siz* double mutants is not suppressed although amounts of N-cadherins are reduced. Therefore, we overexpressed Schizo in photoreceptor cells and performed gene dose experiments and found that only the reduction of the *abi* suppressed the Schizo-induced overexpression phenotype.

In *Salmonella* host cell invasion, the Arf6 GTPase was shown to affect indirectly actin polymerization by activating the GEF ARNO (Humphreys et al., 2013). Arf6 is activated by EFA6 or BRAG and recruits the autoinhibited GEF ARNO to the plasma membrane. The autoinhibition of ARNO is released by the binding of activated Arf6 to its PH domain. As a consequence, ARNO activates Arf1, which induces together with Rac1 the activation of the WAVE Regulatory Complex (Koronakis et al., 2011; Humphreys et al., 2013). Since we found no evidence that *scar/wave* or *wasp* exert a regulatory influence on Schizo, we propose that Abi might be part of a dosage-dependent regulatory feedback mechanism following Arp2/3-dependent actin polymerization.

The data presented in this study point towards a novel model for the regulation of the Arf1 GEF Schizo (Fig. 6). First, we suggest that amounts of N-cadherin are regulated by the Sec7 GEF Schizo and that the N-terminal region of Schizo is crucial for the regulation of its activity (Fig. 6A). The Abl-interacting partner Abi might compete with N-cadherin for Schizo binding and disturbs the removal of N-cadherin. However, since the expression of the constitutive-active form of UAS-Sec7-PH decreases amounts of N-cadherin in the absence of the N-terminal region, we propose that Abi acts rather as an allosteric modulator by either inhibiting the binding of myristyolated Arf1 (Fig. 6B) or by inhibiting the binding of the Sec7-PH module to the lipid bilayer (Fig. 6C). A second regulatory mechanism to control Schizo activity might involve the nuclear localization sequence within the GEF domain in the absence of the N-terminal region. The data reported in our study will now be valuable for future structural analyses to determine how Abi prevents GEF activity.

**Figure 6.**
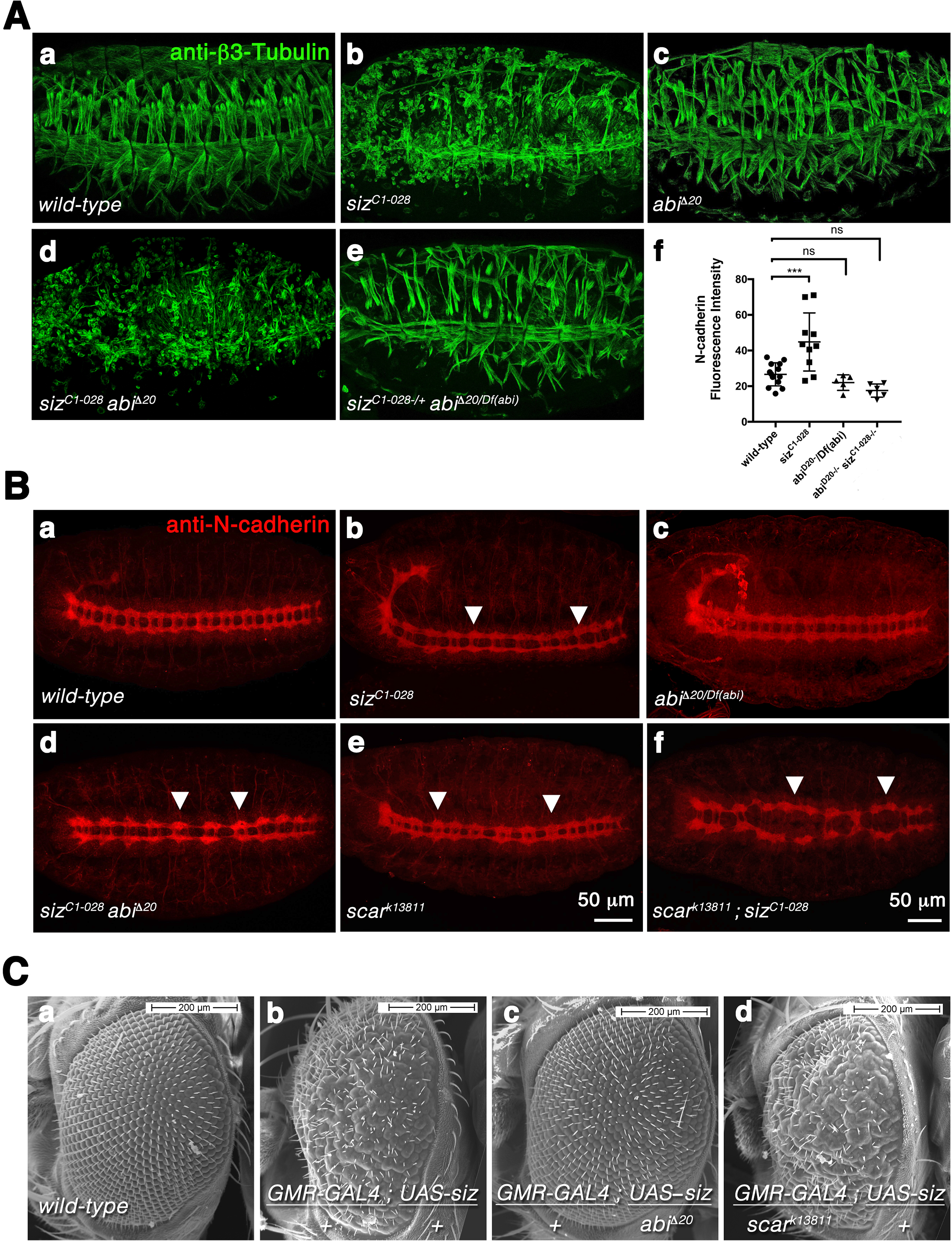
Contingent models indicating how the binding of Abi might antagonize Schizo function. A. The N-terminal region of Schizo interacts with the intracellular region of N-cadherin to regulate amounts of N-cadherin. The activity of Schizo involves the interaction of the Sec7 domain with the Arf1-GTPase and the lipid interaction of the Sec7 and PH domain. The binding of Abi to the N-terminal region of Schizo counteracts N-cadherin regulation in a dosage-dependent manner. B. The binding of Abi to the N-terminal region of Schizo could prevent the binding of the myristoylated Arf-GTPase to Schizo, which inhibits the removal of N-cadherin. C. Alternatively, Abi might prevent the lipid interaction of the Sec7 and PH domain thereby antagonizing Schizo function.

## Acknowledgements

We are grateful to Sven Bogdan for providing Abi full-length, AbiΔSH3 and AbiΔN (Bogdan et al., 2005). We thank Renate Renkawitz-Pohl, Anne Holz and Joanne M. Britto for fruitful discussions and carefully reading the manuscript. We thank Sabina Huhn for technical assistance and Lars Kneifert for perfoming the site-pecific mutagenesis on the *graf* cDNA LD28528 during his bachelor work. Furthermore, we thank the students of the lab2venture project from the Herder school in Gießen that performed a Schizo modifier screen in the *Drosophila* eye and discovered Abi as a Schizo suppressor: Sophie Dönges, Inga Interwies, Anne Mack, Fiona Metsch, Tobias Post and Katrin Stör. This work was supported by the Deutsche Forschungsgemeinschaft (DFG) by the Graduate School GRK 1216 and 2213 as well as by the grant OE311/4-2 to S.F.Ö.

## Material and Methods

### Drosophila stocks and genetics

#### Generation of pUASt-attB-eGFP-siz_1-1313_, pUASt-attB-eGFP-siz_753-1313_ flies and siz-Sec-PH

The *siz_1-1313_* and *siz_753-1313_* construct were amplified from the *LP01489* cDNA by PCR, subcloned into the pENTRTM/D-TOPO^®^ vector and recombined into the Gateway vector pUASt-attB-rfa-eGFP. We used the following primer pairs:

siz_753-1313__f: 5’-CACCATGGAGACGATACGCAAG-3’
siz_753-1313__rev: 5’-TTAGACCTCCGTCGACCGT-3’
siz_Sec7-PH__f: 5’-CACCATGTCGGAGAC-3’
siz_Sec7-PH__rev: 5’-GAGATGGAGTCGTGA-3’

#### graf mutant flies

*y^1^cho^2^v^1^*; *graf* mutants were generated using a single target construct in the CRISPR/Cas9 system as described by Kondo and Ueda (2013). For cloning of the target construct, primers 5’-CTTCGGTCAAAGATCTTATGAGTG-3’ and 5’-AAACCACTCATAAGATCTTTGACC-3’ were used, and the product was ligated to *pBFv-U6*.*2*. The target construct was injected into *y^2^cho^2^v^1^P{nos-phiC31\int*.*NLS}X;attP2(III)*, and established transgenic flies were crossed with *y^2^cho^2^v^1^*;*attP40{nos-Cas9}/CyO* flies for mutagenesis. Twenty-three founder flies were crossed with *y^2^cho^2^v^1^*; *Sco/CyO* flies to establish potential mutants. Genomic regions next to the target sequence of homozygous offspring were analyzed by PCR and screened for deletions.

#### Generation of pUASt-graf-ΔBAR and pUASt-ΔBARΔSH3

The graf-ΔBAR and pUASt-ΔBARΔSH3 constructs were amplified from the full-length *graf* LD28528 cDNA obtained from DGRC. It should be noted that the cDNA clone reported as fully sequenced contains in flybase contains an additional nucleotide at position 467, which leads to a shift of the open reading frame. As a consequence, the deduced protein from this cDNA lacks the BAR domain. To generate full-length Graf, we performed a site-specific mutagenesis on the LD28528 cDNA to remove the additional nucleotide.

#### Other stocks

The hypomorphic *schizo* EMS allele *siz^C1-028^* was generated by Hummel et al. (1999) and the CNS and muscle phenotype of *siz^C1-028^* was characterized and described in Önel et al. (2004) and Dottermusch-Heidel et al. (2012). UAS-*Arf1T31N-eGFP* was generated and described in Dottermusch-Heidel et al. (2012). *abi^Δ20^* mutants were kindly provided by Sven Bogdan (Stephan et al., 2011). The *twist-*GAL4 driver line SG24 was obtained from the Bloomington Stock Center. As *abi* deficiency we used the deficiency line Df(3R)BSC617 (BL25692) from the Bloomington Stock Center. Further GAL4 driver lines that were used in this study are *Mef*-GAL4 (Ranganayakulu et al., 1996) and TGX *twist*-GAL4 (from A. Michelson). As blue balancers we used *Dr*/TM3 *Dfd-lacZ* and *If/*CyO *hg-lacZ*. All crosses were performed at 25°C using standard methods.

### Yeast two-hybrid assay

Yeast two-hybrid experiments were carried out using the Matchmaker GAL4 Two-Hybrid System 3 (Takara Clontech) according to the manufacturer’s instructions. For construct generation, the *schizo* cDNA LP01489 from DGRC was used. The following primers were used to generated the different *schizo* and *graf* constructs:

siz_1-323_ 5’-CATATGATGTCCAGGTGTGA-3’ and 5’-GGATCCTCGCACTCCGC-3’
siz_1-758_ 5’-CATATGATGTCCAGGTGTGA-3’ and 5’-
siz_324-753_ 5’-CATATGATGGCCCGTAACG-3’ and 5’-GGATCCTATCGTCTCCGAC-3’
siz_754-993_ 5’-CATATGATGCGCAAGCGAC-3’ and 5’-GGATCCCACACCAGGTCG-3’
siz_994-1313_ 5’-CATATGATGCATCAGCGCG-3’ and 5’-GGATCCTTAGACCTCCGTC-3’
siz_324-993_ 5’-GAATTCATGGCCCGTAACGCA-3’ and 5’-CTCGAGCACACCAGGTCG-3’ graf 5’-CATATGATGGGCGGCGGCAAAAAT-3’ and 5’-GGATCCTAATGGT GCGGCTTCAAAT-3’
grafΔBAR 5’-CATATGATGTCAACTAAAAAGCCCGAA-3’ and 5’-GGATCCCTAATGGTGCGGCTTCAAAT-3’
grafΔBARΔPH 5’-CATATGCTGGCTCCCGGCA-3’ and 5’-GGATCCCTAATGGTGCGGCTTCAAAT-3’
grafΔBARΔSH3 5’-CATATGATGTCAACTAAAAAGCCCGAA-3’ and 5’-GGATCCGGTGCCCGTTGA-3’
grafΔBARΔPHΔRhoGAP 5’-CATATGAGCGCCGATATCAA-3’ and 5’-GGATCCCTAATGGTGCGGCTTCAAAT-3’

The products were cloned into the pCRII-TOPO vector (Invitrogen). The bait vector pGADT7 was digested with *Nde*I and *BamH*I. *siz* was cloned with EcoRI and XhoI into the pGADT7. *siz* was cloned with *NdeI* and *EcoRI* into the pGADT7.

The pGADT7-T and pBGKT7-p53 pair was used as a positive control. The candidate interaction pairs were co-transformed into yeast strain AH109, and the transformed yeast cells were selected using synthetic dropout (SD/-Leu/-Trp) medium, and then further selected on SD/-Leu/-Trp-His/-Ade selective medium with X-α-Gal (80 mg/L). Results were obtained after 2 days of growth at 30°C.

### Drosophila cell culture

*Drosophila* S2R+ cells were propagated in 1× Schneider’s *Drosophila* medium (Invitrogen) containing 10% fetal bovine serum at 25°C, and transiently transfected described by Kaipa et al. (2013) by using the FuGENE® HD Transfection Reagent (Promega).

### Immunofluorescence

Embryos were fixed and immunohistochemically analyzed as described by Schäfer et al. (2007). The following antibodies were used at the noted dilutions: rat anti-CadN-Ex#8 (Iwai et al., 1997) 1:50 (Developmental Studies Hybridoma Bank), guinea pig anti-β3Tubulin (Buttgereit et al., 1996; Leiss et al., 1988) 1:10,000, rabbit anti-β -Gal 1:5000 (Biotrend), rabbit anti-GFP 1:500 (abcam). Primary antibodies were detected using the fluorescent-labeled antibodies Alexa-Fluor-488, 568- or 647-conjugated anti-guinea pig, anti-rabbit and anti-rat IgG at a dilution of 1:500 (Invitrogen). DNA was stained with Hoechst reagent (5 g/ml; Sigma-Aldrich), and F-actin was stained with Alexa-Fluor-647–phalloidin (1:100, Invitrogen). For all stainings, specimens were embbeded in Fluoromount-G™ (Thermo Fisher Scientific) and observed under a Leica TCS Sp2 or TCS Sp8 confocal microscope.

### Fluorescence Intensity and Statistical Test

Statistical analyses were performed with GraphPad Prism software (PRISM7). For Figures 2, 3, 5 and EV1 significance was determined by one-way ANOVA with comparison.

### Microscopy and image analysis

A Leica TCS Sp2 (Fig. 1B–E”) and TCS SP8 (Fig. 1F–G’, Fig. 2, Fig. 3, Fig. EV1, Fig. 5) confocal microscope was used for fluorescence imaging. The same parameter settings were used to image all samples of the same type. Embryos were embedded in Fluoromount-G™ (Thermo Fisher Scientific) and scanned using the 20x objective with the galvo scanner at 400 Hz. The raw data of the embryo images were analyzed using the FIJI software (Schindelin et a., 2012). A SUM-stack was generated in Fiji and the mean fluorescence intensity of three nearby areas were also measured to analyze the background fluorescence level. The corrected total relative intensity of each embryo was calculated as follows:

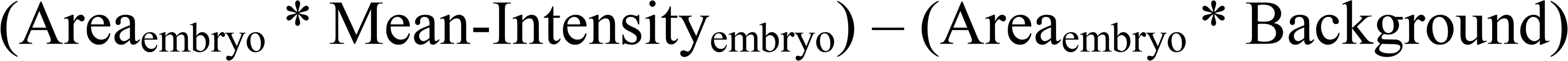

For time-lapse imaging, the Spinning disc microscope from Zeiss was used with the 63x objective.

### Scanning electron microscopy

All *GMR-*GAL4>UAS-*siz*-expressing flies were raised at 25°C. Eyes were fixed in 6% glutaraldehyde and 1% formaldehyde in 0.2 M Hepes buffer for 16 h. Samples were then dehydrated in a 25%, 50%, 70% and 96% ethanol series for 12 h each and finally transferred into acetone by three 10-min changes with 100% acetone. The samples were critical-point-dried by using a Polaron E 3000 (Balzers Union). Samples were attached to sample stubs (Plano GmbH) and sputtered with gold under vacuum using a sputter coater (Balzers Union, Lichtenstein). Scanning electron micrographs of adult fly eyes were taken using a Hitachi S-530 SEM.

## Author contributions

SFÖ initiated, conceived and supervised the project. SL performed and analyzed the experiments in *Drosophila* and performed the yeast two-hybrid assay with Schizo and Abi. CB performed the yeast two-hybrid assay with Graf, Schizo and N-cadherin. CB generated the Siz_753-1016_ construct and performed subcellular localization studies of Schizo-fl and Siz_753-1016_. KHR performed scanning electron microscopy. LC injected the construct to generate *graf* mutant flies by CRISPR/Cas9. SFÖ wrote the manuscript.

## Conflict of interest

The authors declare that they have no conflict of interest.

---

**Figure EV1.**
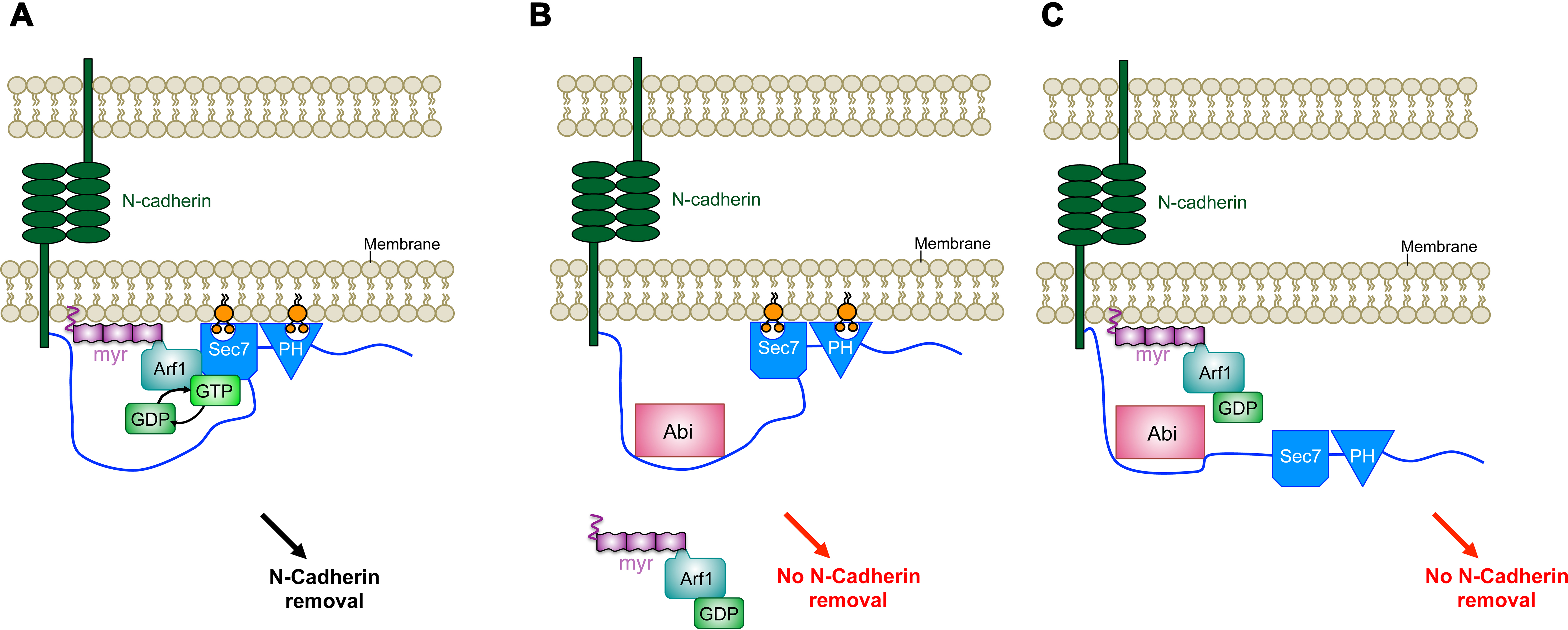
The *Drosophila* RhoGAP protein Graf-1 interacts with the intracellular domain of N-cadherin, but *graf-1* mutants show no defects in myoblast fusion. A. Schematic diagram of full-length and truncated forms of Graf-1 tested for interaction with the intracellular domain of N-cadherin. B,C. Transfected *Drosophila* S2R+ with UAS-*graf-eGFP* or *UAS-graf-mcherry* and stained with DAPi (blue) and Phalloidin (grey). Cells were plated on concanavalin A coated cover slips (B) and on polylysine coated cover slips (C). Scale bar in B 10 μm and scale bar in C 5 μm. D,E. Transfected *Drosophila* S2R+ with *UAS-graf-mcherry* and UAS-*siz-eGFP* (D) or GTP-bound UAS-*Arf1Q71L* (E). Cells were plated on polylysine coated cover slips and stained with DAPI (blue). Both proteins partially colocalize with Graf-mcherry. Scale bars 5 μm. F. Protein structure of Graf-1. The exon-intron structure of the BAR domain is highlighted underneath. The 18-bp target sequence of the gRNA is indicated in red, adjacent to NGG protospacer adjacent motif (PAM) sequence in green. Obtained mutants were analyzed by PCR. Red dashes indicate the identified mutations. G,H. Lateral view of stage 16 embryos stained with anti-β3-Tubulin. (G) Homozygous *grafΔ20* mutant embryos. (H) Transheterozygous *grafΔ20/grafΔ4* mutant embryo. I. Quantification of the amounts of N-cadherin in wild-type, homozygous *siz^C1-028^* mutant embryos, embryos expressing UAS-*grafΔBAR-eGFP and* UAS-*grafΔBARΔSH3-eGFP* with *Mef-*GAL4. The total fluorescence intensity of 6 to 13 embryos was measured for each experiment. *Mef*-GAL4≫UAS-*grafΔBAR-eGFP* and *Mef*-GAL4≫ UAS-*grafΔBARΔSH3-eGFP* embryos show no significant increase in N-cadherin amounts. Bars represent mean ± s.d. P-values were calculated using the Dunnett’s multiple comparison test. p^**^< 0,0032 compared to wild-type. ns = not significant.

